# Unique B Cell and Germinal Center Responses in Mice with Severe versus Mild *Orientia tsutsugamushi* Infection

**DOI:** 10.1101/2025.03.27.645674

**Authors:** Casey Gonzales, Yuejin Liang, Joseph Thiriot, Hui Wang, Dario Villacreses, Jiaren Sun, Lynn Soong

## Abstract

The intracellular bacterium *Orientia tsutsugamushi* (*Ot*) is the causative agent of scrub typhus, an emerging and neglected tropical disease. Immunity against *Ot* infection in patients is known to be short-lived, as circulating antibodies can wane around one-year post-infection. However, its underlying mechanisms remain unclear; host immune signatures to clinically prevalent *Ot* strains are also undefined. In this study, we infected C57BL/6 mice with two clinically prevalent *Ot* strains with distinct virulence to examine splenic B cell and germinal center (GC) responses. While highly virulent Karp strain-infected mice had 50% mortality rates, intermediate virulent strain Gilliam-infected mice survived, gained body weight, and maintained relatively low bacterial burden levels during infection. Yet, Gilliam infection resulted in strong splenic B cell responses, as judged by absolute cell numbers of total splenic B cells, follicular B cells, and marginal zone (MZ) B cells alongside with significantly higher titers of total IgG and IgM antibodies at the late stage of infection. While altered splenic architecture with increased distance between white pulp regions was detectable in both strains, GC disorganization/collapse and MZ abrogation were only detected in Karp-infected spleens. To define the driving force of GC loss, we compared RNAseq profiles between spleens infected with the Karp or Gilliam strain, and found significantly upregulated inflammation biomarkers (*Ccl2*, *Il33*, *Ifng*) and severe scrub typhus-related inflammatory pathways (*Il1*, *Il6*, *Tnf*, and *Ifng*) in Karp infection at day 4 (D4, early stage of infection). IPA analysis further showed upregulated neutrophil degranulation and defense response pathways in Karp-infected spleens, while Gilliam-infected spleens showed upregulation of phagocytosis signaling pathways. Flow cytometry confirmed a marked influx of activated myeloid cell subsets (neutrophils, M1 macrophages, and inflammatory monocytes) in the spleen during Karp infection as compared to Gilliam infection, indicating excessive inflammatory infiltration in highly virulent *Ot* strain infection. Collectively, our findings establish several *Ot* strain-related immune signatures which help fill the knowledge gap of deficiencies in the host adaptive immune response during acute scrub typhus.

**Author Summary:** The intracellular bacterium *O. tsutsugamushi* (*Ot*) is the etiologic agent of the neglected tropical disease scrub typhus. It is well-known that immunity against *Ot* infection is short-lived, yet no in-depth studies have examined or compared adaptive immunity or deficiencies at cellular and molecular levels during acute *Ot* infection with clinically relevant *Ot* strains. In this study, we used two clinically prevalent *Ot* strains that cause differential diseases outcomes in the C57BL/6 murine model to examine B cell and germinal center (GC) responses. In sharp contrast to Karp infection, which caused 50% mortality rates by D12, Gilliam infection elicited self-limiting infection with low tissue splenic bacterial burdens. Yet, Gilliam-infected mice demonstrated robust humoral immune responses, as demonstrated by strong IgM and IgG antibody titers at D12 and absolute numbers of splenic B cell subsets at D8 and D12. While both *Ot* strains altered typical splenic white pulp architecture, only Karp infection abrogated GC and MZ structures. Comparison of RNAseq profiles of Karp-vs. Gilliam-infected spleens revealed several unique features, including Karp infection-associated upregulation of inflammation genes (*Ccl2*, *Il33*, *Ifng*) and inflammatory pathways (*Il1*, *Il6*, *Tnf*, *Ifng*) at D4, as well as upregulated leukocyte recruitment, neutrophil degranulation, and defense response pathways. In sharp contrast, Gilliam infection upregulated pathways involved in phagocytosis immunoregulatory interactions. Flow cytometry confirmed significant influx and activation of myeloid cell subsets (neutrophils, macrophages, M1 macrophages, monocytes) during Karp infection. Our study reveals unique patterns in the differential immune responses to two clinically prevalent *Ot* strains and helps understand potential mechanisms of immune alterations during severe scrub typhus.

## Introduction

*Orientia tsutsugamushi* (*Ot*) is an obligately intracellular bacterium and causative agent of the understudied and underdiagnosed febrile illness, scrub typhus. Within endemic regions, commonly known as the tsutsugamushi triangle, there are an estimated one million cases annually, although the actual number of cases is likely much higher than this estimation due to lack of surveillance, rural prevalence, and difficulties with diagnosis [1]. Scrub typhus can be life-threatening, particularly in cases with delayed or inadequate antibiotic treatment, where median case fatality rates increase from 1.4% with treatment to 6% without treatment [2–4]. In severe scrub typhus cases, multiple organs can become infected and manifestations such as acute respiratory distress syndrome, acute renal failure, meningoencephalitis, disseminated intravascular coagulation, or septic shock can occur [5]. Although historically confined to endemic regions, the recent reports of new *Orientia candidatus* species in Chile (*O. chiloensis*) and Dubai (*O. chuto*), along with the detection of *Ot* in mite vectors in North Carolina, has drawn renewed attention to this neglected yet life-threatening tropical disease [6–9].

At least 20 antigenically distinct *Ot* strains have been reported [10]. These strains show a relationship between their epidemiological prevalence and geographic distribution. For example, isolates related to the prototype Karp, Gilliam, and Kato strains are found throughout the tsutsugamushi triangle, especially in East Asian countries such as Japan, South Korea, and China [10, 11]. To this effect, Karp and Gilliam strains are clinically relevant, representing approximately 65% and 26% of global cases, respectively [12]. In humans, these strains can cause symptomatic scrub typhus. A recent study in Guangzhou showed that patients with Karp infection presented with greater disease severity than patients with Gilliam infection, exemplified by higher likelihood of hospitalization and longer hospital stay, higher risk of multiorgan involvement and dysfunction, longer fever duration, and greater intensive care unit admission [13]. Furthermore, it has also been shown that Karp-infected patients have significantly higher bacterial burdens in their blood than Gilliam-infected patients [14].

It is well-known that immunity to scrub typhus is nondurable, waning as early as one year post-infection in some scrub typhus patients, with reports of heterologous immunity vanishing as early as one month post-infection [15–17]. Cellular immunity, which is critical to control acute *Ot* infection, declines relatively quickly after infection. A study by Ha *et al* identified that type specific antigen 56 (TSA56)-specific CD4^+^ and CD8^+^ T cells in scrub typhus patients declined only one year after infection and were almost undetectable by two years post-infection [16]. Humoral immunity follows a similar trend, as 50% of *Ot*-infected individuals were shown to have reverted to sero-negativity only 49 weeks post-infection [15]. Another study demonstrated that antibodies to ScaA and TSA56 had returned to baseline by two years-post-infection, with antigen-specific ScaA antibodies declining quicker than those against TSA56 [16]. Beyond the relatively short-lived cellular and humoral immunity after initial infection, there are additional difficulties with immunity after infection. Heterologous cross protection between strains is weak due to the antigenic differences among strains [18]. In some patients challenged with heterologous strains led to bacteremia only one month after initial infection, and by one year-post-infection, heterologous challenge led to scrub typhus in all human volunteers [17]. During acute immune responses, scrub typhus patients display inflammatory cytokine profiles, with IFNγ, TNFα, IL-6, IL-8, and IL-12p40 showing an association of type 1 responses with *Ot* infection [19–22]. Scrub typhus-associated chemokines include MCP-1, MIP-1β, and CXCL10 [19, 20]. Lastly, the cytokines TNFα, IL-8, and IL-10 show positive correlation with severe scrub typhus patients [20–22].

Experimental murine models are valuable tools to characterize *Ot* strain virulence and immunity. Karp is categorized as being high virulence and Gilliam as being intermediate virulence [23, 24]. For example, when CD-1 outbred mice (a highly susceptible scrub typhus model) were infected with the same viable dose of Karp or Gilliam strain, Karp infection caused 100% lethality, while Gilliam infection caused only 50% [24]. Most importantly, Karp infection resulted in higher bacterial burdens in various organs and extremely robust inflammatory response as compared to Gilliam infection, exemplifying the diverse immune signatures induced by different *Ot* strains [24]. Beyond the acute response to infection, Mendell *et al.* demonstrated that Karp heterologously infected C57BL/6 mice showed diminishing immunity only 9 months after initial Gilliam infection, establishing that cross-protective, heterologous immunity is short-lived [25].

While patient-based studies differential immune responses to distinct *Ot* strains are lacking, animal model-based and *in vitro* studies have offered valuable information. For example, our group recently reported that Karp, but not Gilliam strain, could cause severe disease in C57BL/6 mice [24]. Compared to Gilliam-infected lung tissues, Karp-infected tissues had significantly higher infiltration of innate immune cells (monocytes, M1 macrophages, activated neutrophils, and activated NK cells), increased pathological lesions, and elevated expression of proinflammatory genes like *Ccl2/3/4/5* and *Ifng* [24]. Similarly, a more recent study showed that highly virulent strains of *Ot* were associated with elevated serum levels of MCP-1, IL-6, IL-10 and IFNγ in infected mice [26]. Proinflammatory genes *Tnf*, *Cxcl1*, *Ccl2*, and *Ccl5* were more highly expressed in Karp-vs. Gilliam-infected bone marrow-derived macrophages [24].

Most scrub typhus-focused immunological studies are done in target organs (such as the lungs) and target cells (macrophages, endothelial cells, fibroblasts). In 2023, our group reported the first evidence that severe Karp infection induced the loss of GCs, disorganization of splenic microarchitecture, and attenuation of B cell response pathways [27]. In general, GC collapse is associated with poor immunity, as the GC response is critical to the development of humoral immunity after infection [28]. It is therefore not surprising that some pathogens have evolved diverse strategies to manipulate host B cell responses to subvert the humoral immune responses, including the disorganization of splenic microarchitecture [29, 30]. Yet, no studies have defined whether *Ot* virulence can affect the development of adaptive immune responses. Given the striking differences in disease outcomes and cellular response following Karp vs. Gilliam infection in outbred and inbred mouse models, analysis of how splenic immune responses are altered due to the virulence range of *Ot* strains is essential.

In this study, we used our established C57BL/6 models to examine spleen at whole tissue, cellular, and molecular levels to assess whether mice with severe scrub typhus (Karp strain) or self-limited infection (Gilliam strain) had differential effects on splenic architectural organization and GC formation, and whether these strains elicited unique immune responses in these immunological niches that correlated with disease pathogenesis. In contrast to Karp-infected animals, Gilliam-infected mice had minimal bacterial growth in the spleen during acute infection stages, but they developed higher sera IgM and IgG antibody responses at D12, as well as greater induction of B cell, follicular B cell, and MZ B cell numbers. We found that Karp infection exclusively elicited GC collapse and disorganization, as well as splenic MZ abrogation, as judged by immunohistology and confocal microscopy. On one hand, Karp-associated transcriptomic fingerprints revealed significant induction of inflammation genes (*Ccl2*, *Il6*, *Il10*, *Ifng*) and related pathways (TNF, IFNγ, IL-1, IL-17), as well as innate immune cell influx (monocytes, macrophages, neutrophils) at D4 and D8. Flow cytometric analysis confirmed Karp infection-associated increase of influx and activation of myeloid cell subsets, in contrast to Gilliam infection-associated expression of adaptive immune responses. This is the first comprehensive study to evaluate differential immune responses to two prevalent *Ot* strains in the context of the splenic immunological niche, revealing new insights into the collapse of GCs and the virulence-associated alterations in adaptive immune responses.

## Materials and Methods

### Mouse infection and ethics statement

Female C57BL/6 mice (#000664, Jackson Laboratory) were maintained under specific pathogen-free conditions. Animals used were at 8-12 weeks of age, following protocols approved by the Institutional Animal Care and Use Committee (IACUC#1902006 and #2101001A) at the University of Texas Medical Branch (UTMB) in Galveston, TX. Mouse infection studies were conducted in the Galveston National Laboratory ABSL3 facilities using procedures approved by the Institutional Biosafety Committee, in accordance with Guidelines for Biosafety in Microbiological and Biomedical Laboratories. UTMB complies with the USDA Animal Welfare Act (Public Law 89-544), the Health Research Extension Act of 1985 (Public Law 99-158), the Public Health Service Policy on Humane Care and Use of Laboratory Animals, and the NAS Guide for the Care and Use of Laboratory Animals (ISBN-13). UTMB is registered as a Research Facility under the Animal Welfare Act and has current assurance on file with the Office of Laboratory Animal Welfare, in compliance with NIH policy. Both *Ot* Karp and Gilliam stocks were prepared in L929 cells, and the inoculum infectivity titer of prepared stocks was determined, as in our previous report [31]. Mice were intravenously inoculated with a viable dose of (6.8 × 10^4^ FFU, 200 µl) of Karp, Gilliam, or PBS (mock) and monitored daily for weight loss, signs of disease, and survival. Disease scores were assigned to mice daily, ranging from 0-5, based on an institute approved protocol for animal sickness [32, 33]. A score of 0 represented normal behavior; 1 represented <5% weight loss with normal activity; 2 represented 6-10% weight loss with ruffled fur between shoulders; 3 represented 11-19% weight loss with pronounced ruffled fur, hunched posture, erythema, and reduction in food/water taken; 4 represented 20-25% weight loss with reduced activity, bilateral conjunctivitis, minimal food/water intake; 5 represented greater than 25% weight loss or non-responsiveness and the animal was humanely euthanized. At D4, D8, and D12 post-infection, serum and spleen tissue samples (4-5/group) were collected and prepared for the subsequent immunological analysis. All independent studies were performed with the same bacterial stock. Data shown are representative of two independent experiments.

### Flow cytometry

Spleens were passed through 70-μm cell strainers in RPMI 1640 medium to prepare single-cell suspensions which were then treated with Red Blood Cell Lysis Buffer (Sigma-Aldrich). Cells were first blocked with FcγR blocker (BioLegend), followed by staining with Fixable Viability Dye eFluor 780 (eBioscience) or LIVE/DEAD Fixable Blue Dead Cell Stain (Thermo Fisher Scientific) and fluorochrome-labeled antibodies (Abs). The Abs below were purchased from either BD Biosciences, BioLegend, Invitrogen, or Tonbo Biosciences for B and T cell staining: BV785-anti-B220, V450-anti-CD3, APC-anti-CD138, PerCPCy5.5-anti-CD38, Alexa fluor488-anti-GL7, PE-Dazzle594-anti-CD23, PE-Cy7-anti-CD21/35, PE-anti-IgM, BV510-anti-IgD, PE-Cy7-anti-CD3, PerCP-Cy5.5-anti-CD4, FITC-anti-CD8, BV711-anti-CD44, BV605-anti-PD-1, BV421-anti-CXCR5, and PE-anti-FOXP3. For myeloid cell staining, the following Abs were purchased from BD Biosciences, BioLegend, and eBioscience: FITC-anti-CD11b, APC-anti-Ly6G, PE-Dazzle-594-anti-Ly6C, RB545-anti-CD80, APC-Cy7-anti-CD63, BV421-anti-F4/80, RB780-anti-SCA-1, and BV650-anti-CCR2. Cells were fixed in 2% paraformaldehyde overnight at 4°C prior to data acquisition. Data were acquired on a BD FACSymphony A5 in the UTMB Flow Cytometry Core and analyzed by using FlowJo software version 10.7.2 (BD Bioscience).

### Immunohistology

Spleen tissues were fixed in 4% paraformaldehyde and 5% sucrose/PBS overnight. Tissues were transferred into 20% sucrose/PBS for 24 h at 4°C, followed by 30% sucrose/PBS for another 24 h at 4°C. Spleens were embedded in O.C.T. compound (Sakura Finetek). Frozen cryosections (7-μm) were blocked with 1% BSA and 0.3 M glycine in PBS for 30 min. They were then incubated with rat IgG2a anti-B220 (1:175), rat IgM anti-GL-7 (1:175), biotin anti-CD3 (1:200) Abs for 1 h at room temperature (BioLegend, clones RA3-6B2, GL-7, 17A2, respectively). Cryosections were then stained with secondary Abs, Alexa Fluor 594-conjugated mouse anti-rat IgG2a (1:200, clone MRG2a-83, BioLegend), Alexa Fluor 488-conjugated goat anti-rat IgM (1:200, clone A21212, Invitrogen), streptavidin cyanine 5 (1:200, BioLegend) for 1 h. Staining with secondary antibodies and primary antibodies alone served as negative controls. For each section, at least 4-5 fields of each spleen section were imaged at the UTMB Optical Microscopy Core on a Zeiss LSM 880 confocal microscope (Carl Zeiss Microscopy LLC) equipped with ApoTome and Zen imaging software. The 488, 561, and 633 excitation lasers under 63x oil immersion objective were used. Acquisition settings were identical among samples of different experimental groups, and representative images are presented from each time point.

### Quantitative reverse transcription PCR (qRT-PCR)

Splenic tissues were collected and incubated in RNA*Later* (Qiagen) at 4°C overnight for inactivation. Tissues were homogenized in a BeadBlaster 24 Microtube Homogenizer (Benchmark Scientific) with RLT lysis buffer (Qiagen) and metal beads. Total RNA extraction was done using RNeasy Mini Kit (Qiagen), and the cDNA synthesis was completed utilizing iScript Reverse Transcription kit (Bio-Rad). cDNA was synthesized in a 10 μL reaction mixture containing 5 μL of iTaq SYBR Green Supermix (Bio-Rad) and 0.5 μM each of gene-specific forward and reverse primers. qRT-PCR assays were performed on a CFX96 Touch Real-Time PCR Detection System (Bio-Rad), and PCR assays were denatured for 30 seconds at 95°C, followed by 40 cycles of 15 seconds at 95°C, and 60 seconds at 60°C. Melt curve analysis was performed to check specificity of amplification. Relative quantitation of mRNA expression was calculated utilizing the 2^-ΔΔCT^ method. Primers used in qRT-PCR analysis are listed in S1 Table.

### Quantitative PCR for tissue bacterial burden

DNA extraction was performed by using a DNeasy Blood & Tissue Kit (Qiagen) followed by qPCR assay, as in our previous report [34]. The copy number for the 47-kDa gene was determined by known concentrations of a control plasmid containing single-copy insert of the gene. Gene copy numbers were determined via 10-fold serial dilution of the *Ot* 47-kDa plasmid. Bacterial burdens were normalized to total nanogram (ng) of DNA per µL for each of the same samples. Data are expressed as the gene copy number of 47-kDa protein per ng of DNA.

### Enzyme-linked immunosorbent assay (ELISA)

Serum was separated from whole blood samples in blood separation tubes (BD Bioscience) by centrifuging at 9000 g for 2 min. For analysis of antigen-specific Ab responses, 96-well plates were coated with 2 μg/mL of recombinant Karp TSA56 and 2 μg/mL of recombinant Gilliam TSA56 (Karp TSA56 and Gilliam TSA56 mixed at a 1:1 ratio, recombinant proteins were generated by Genscript) in PBS and blocked with 0.5% BSA. Serum was diluted 1:3 until endpoint titers were determined. Detection was performed utilizing the following horseradish peroxidase-conjugated primary antibodies that were diluted 1:3000 in blocking buffer: goat anti-mouse IgM (1021-05, Southern Biotech) and goat anti-mouse IgG (1030-05, Southern Biotech). Visualizing reagent utilized was the 1-Step Ultra TMB ELISA Substrate Solution (Thermo Fisher Scientific). Optical density was measured on the BioTek Epoch microplate spectrophotometer. Area under the curve (AUC) analysis was performed on each curve (n=5) at every timepoint [35].

### Splenic RNAseq

Splenic tissues were incubated in RNA*Later* (Qiagen) at 4°C overnight, followed by RNA extraction utilizing the RNeasy Mini Kit (Qiagen). RNAseq analysis was performed by Novogene (San Jose, CA). Following quality control assessment, mRNA was purified from total RNA with poly-T oligo-attached magnetic beads. Fragmentation, cDNA synthesis and library preparation were then performed. The Illumina Novaseq Platform NovaSeq X Plus was used for sequencing. The *Mus musculus* genome build mm10 was used as a reference genome. Raw data were first evaluated for quality control to exclude reads with adapter, poly-N, and low quality, and thus only clean data were further examined. Normalization and analysis were completed by Novogene using featureCounts v1.5.0-p3 for quantification of gene expression levels, DESeq2 R package and for differential expression analysis, and clusterProfiler R package for Gene Ontology, KEGG, Reactome, Wikipathways, and Pathway Interaction DB databases enrichment analysis of differentially expressed genes. Ingenuity Pathway Analysis (IPA) was performed on Karp versus Gilliam comparisons at each time point to identify canonical pathways with results filtered by a Benjamini-Hochberg corrected *p*-value. Results were reported with z-scores and filtered by an adjusted *p*-value < 0.05. All RNAseq data discussed in this publication have been deposited in NCBI’s Gene Expression Omnibus and are accessible through GEO Series accession number GSE290855 (https://www.ncbi.nlm.nih.gov/geo/query/acc.cgi?acc=GSE290855).

### Statistical analysis

All data, excluding bacterial RNAseq data, were analyzed by using GraphPad Prism software. Data were presented as mean ± standard deviation (SD) or standard error of mean (SEM). Bodyweight change data were statistically analyzed with two-way ANOVA and Šídák’s multiple comparisons test. Survival data were assessed using a survival curve comparison and Log-rank (Mantel-Cox) test, log-rank test for trend, and Gehan-Breslow-Wilcoxon test. Disease score and splenic bacterial burden qPCR data were analyzed with one-way ANOVA and Tukey’s multiple comparisons test Post Hoc for comparisons between groups. ELISA AUC data, RT-PCR, and flow cytometry data were analyzed with one-way ANOVA and Tukey’s multiple comparisons test Post Hoc for comparisons between groups. Statistically significant values are denoted as \**p* < 0.05, ** *p* < 0.01, *** *p* < 0.001, and **** *p* < 0.0001, respectively, or ns for no significance.

## Results

### Distinct disease outcomes induced by *Ot* Karp vs. Gilliam strain in mice

Karp and Gilliam strains account for most of human scrub typhus, representing approximately 65% and 26% of global cases, respectively [12]. Yet, no detailed studies have compared the development of acute splenic adaptive immune responses elicited by infection with the Karp or Gilliam strain. Here, we intravenously inoculated C57BL/6 mice with an equivalent viable dose of Karp or Gilliam and measured kinetics of disease progression each day. We found that Karp-infected mice exhibited body weight loss beginning at D4 and reaching more than 20% body weight loss at D10, while Gilliam-infected mice lost no weight during infection (Fig 1A). Concurrently, Karp-infected mice began to succumb to infection at D10, leading to 50% lethality at D12 (Fig 1B). Conversely, Gilliam-infected counterparts demonstrated 100% survival (Fig. 1B). Karp-infected mice also displayed significantly higher disease scores from D7 through D12, reaching a peak score at D10 (Fig 1C); however, Gilliam-infected counterparts showed no signs of disease. We also found significantly increased splenic bacterial burdens in Karp infection at D4 and D8 (Fig 1D), resulting in a 15-fold difference in bacterial burdens between the two strains at the peak of infection (D8). Overall, we demonstrated that for the inoculation dose we used, Karp strain caused a severe and sublethal infection, while Gilliam strain caused a self-limiting infection with no signs of disease. These results are in line with other studies utilizing the Karp and Gilliam strains in inbred and outbred murine models, respectively [24, 25].

**Fig 1.**
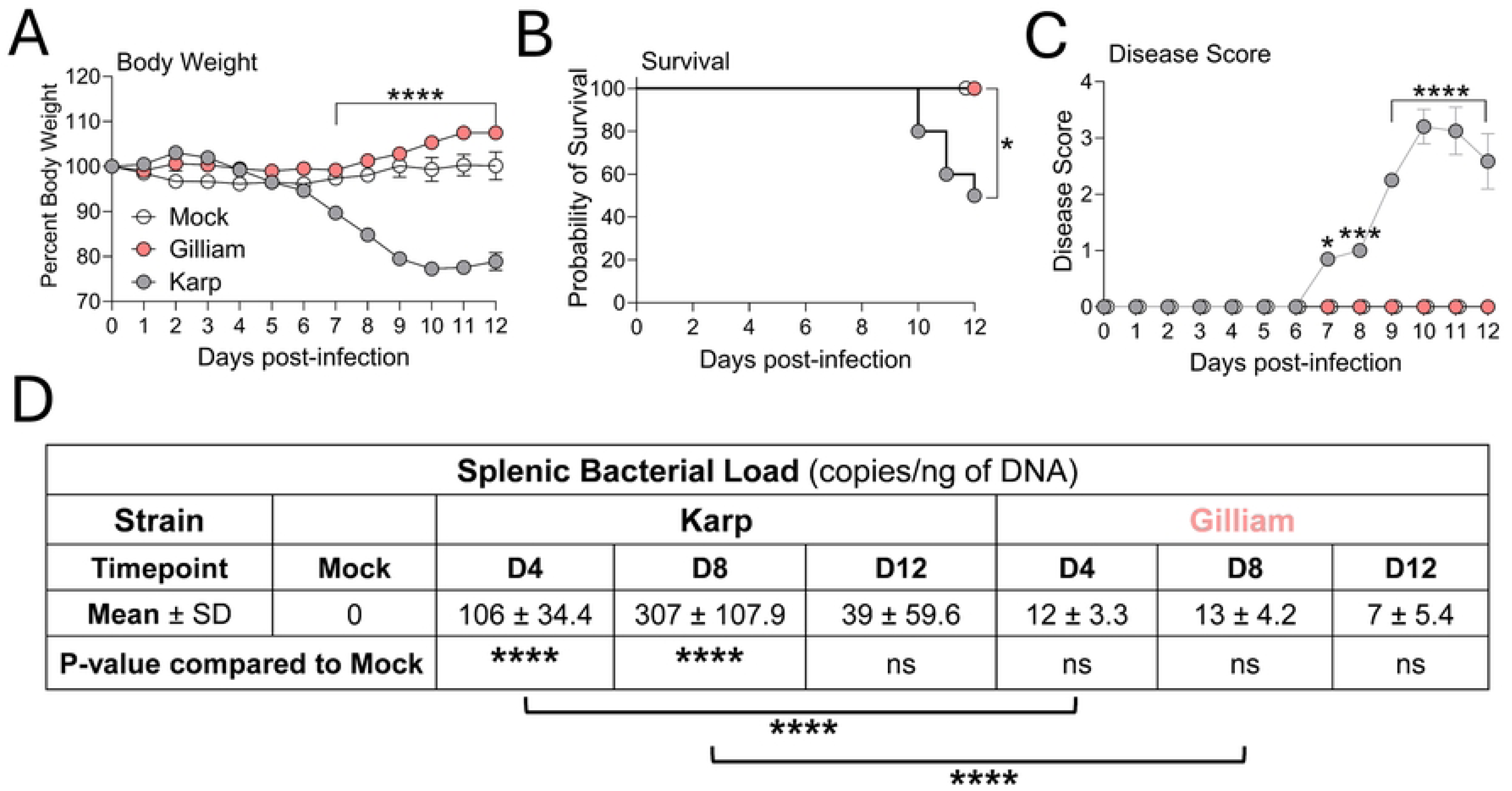
Disease progression kinetics in *Ot* Karp-or Gilliam-infected mice. C57BL/6 mice (5 per group) were intravenously infected with Karp or Gilliam strain (6.8 × 10^4^ FFU) or PBS as an uninfected mock control and monitored daily for body weight (A) and survival (B). PBS was used as a mock control. C) Disease score, ranging 0-5, was observed daily. Mice (n = 5 per group) were euthanized at D0 (Mock), D4, D8, and D12, and spleens and sera were harvested. D) Splenic bacterial burden was measured by qPCR. Data are shown as mean ± SD from a single experiment and are representative of two independent experiments with similar trends. For statistical analysis, two-way ANOVA was used with a Šídák’s multiple comparisons test (A). Survival curves were analyzed by survival curve comparison and Log-rank (Mantel-Cox) test, logrank test for trend, and Gehan-Breslow-Wilcoxon test (B). Disease score and splenic bacterial load data were analyzed by one-way ANOVA and Tukey’s multiple comparisons test (C, D). Asterisks or ns were representative of comparison between Karp- and Gilliam-infected mice at each timepoint, unless otherwise stated. ns, *, *p* <0.05; **, *p* <0.01; ***, *p* <0.001; ****, *p* <0.0001.

### Antigen-specific IgM and IgG antibody responses triggered by Karp vs. Gilliam infection

We previously reported that lethal Karp infection induced serum IgM antibody titers at D4, followed by significant increases in antibody titers of IgG subtypes (particularly IgG2c) at D8 [27]. To test whether antibody levels differed between the two disease models, we measured serum total IgM (Fig 2A) and IgG (Fig 2B) titers via antigen-specific ELISA and then performed area under the curve (AUC) analyses. In comparison to the mock group (background), both Karp and Gilliam-infected mice had elevated and comparable IgM levels at D4 and D8; however, IgM titers displayed differential trends at D12, as IgM titers in Karp-infected mice were significantly lower (approximately 3.7-fold) than those seen in Gilliam infection. In terms of IgG titer and AUC analyses, D4 and D8 of Karp infection respectively stimulated 3- and 2.5-fold higher responses than those of Gilliam infection. By D12, however, Gilliam infection induced significantly greater (1.6-fold higher) IgG responses than Karp infection. It is generally believed that early IgM, or even IgG responses are derived from the extrafollicular or B1 B cell response [30, 36]. Early IgG responses in Karp infection are likely derived from extrafollicular responses, which are likely driven by the Th1-skewed cytokine profile associated with scrub typhus [27]. However, higher IgM and IgG antibody responses seen in Gilliam-infected mice at D12 may be attributed to GC responses [36].

**Fig 2.**
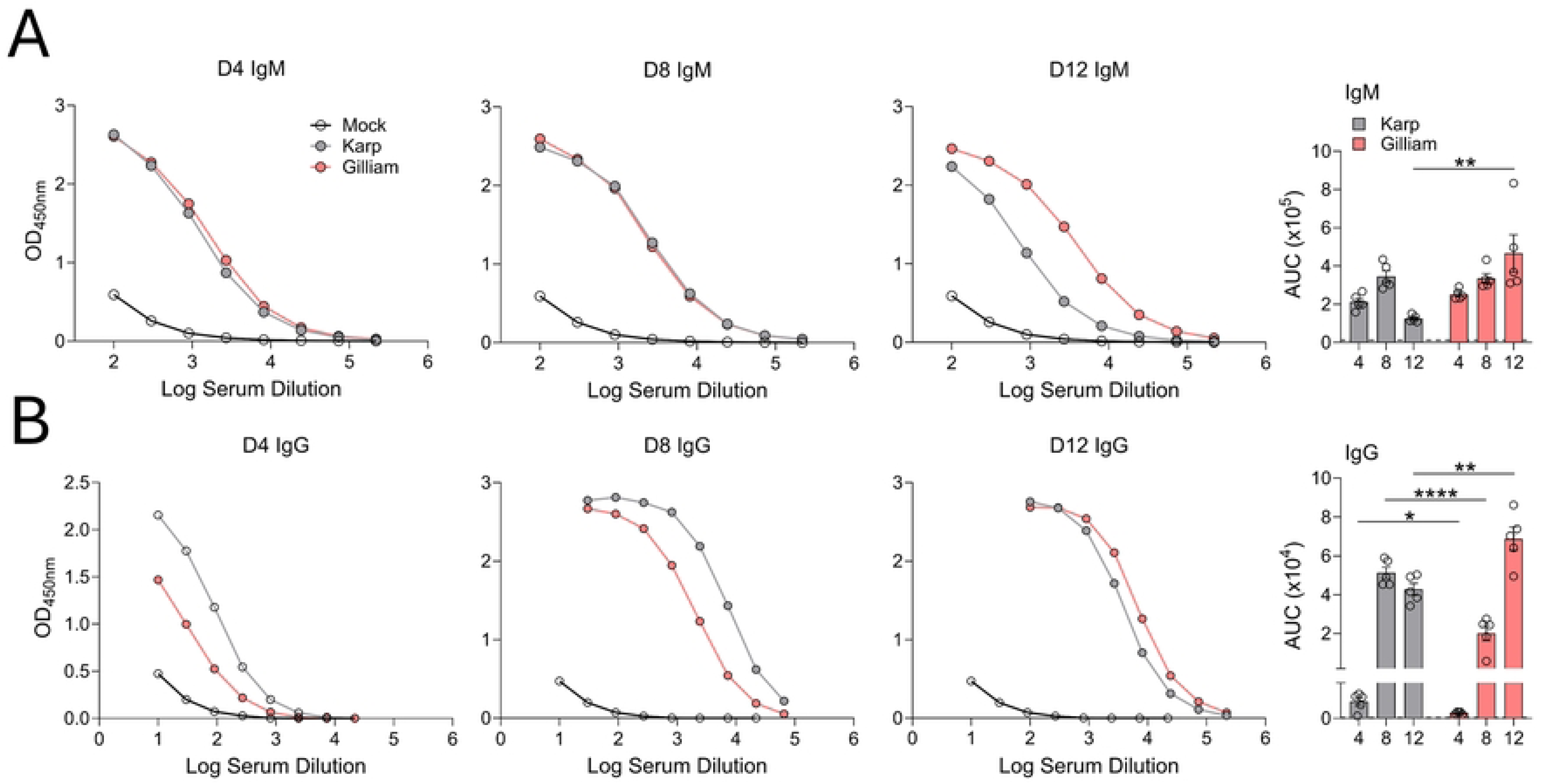
Differential serum IgM and IgG antibody response in Karp-vs. Gilliam-infected mice. Mice were infected, as described in Fig. 1. A) Serum antibody titers specific to recombinant TSA56 derived from Karp and Gilliam sequences were measured in Karp- and Gilliam-infected mice and mock mice (5 per group) via indirect ELISA assay. A) ELISA curves for IgM responses and B) IgG responses are shown. C) AUC analysis of ELISA antibody curves was performed on IgM and IgG data. Data are shown as mean ± SEM from a single experiment and are representative of two independent experiments with similar trends. For statistical analysis, one-way ANOVA was used with a Tukey’s multiple comparisons test for comparison of mock-, Karp-, and Gilliam-infected samples at each timepoint. Asterisks were representative of comparison between Karp- and Gilliam-infected mice at each timepoint. *, *p* <0.05; **, *p* <0.01; ***, *p* <0.001; ****, *p* <0.0001.

### Severe splenic GC disorganization during Karp infection

Our previous report with severe Karp infection (100% lethality) revealed significant alterations in splenic architecture as well as GC collapse [27]. To investigate if such disorganization would also occur in a model with 50% lethality or self-limiting infection, we performed immunostaining of splenic B cells, T cells, and GC B cells. Overview scans revealed comparable and progressive changes of white pulp regions in both infection groups (Figs 3A and 3B). At D4, Karp- and Gilliam-infected spleens showed well-organized white pulp regions, including expanded T cell zones and discernable GCs within B cell follicles (Figs 3A and 3B). At D8, spleens from Karp- and Gilliam-infected mice began showing alterations inside and outside of white pulp structures, including reduced size of T cell zones and an increase of T cell frequency outside of white pulp regions. By D12, splenic architecture was significantly changed compared to D4, with increased space between white pulp regions and T cells outside of T cell zones in both Karp- and Gilliam-infected mice. While these changes seemed to be generally consistent regardless of *Ot* strains used, Karp-infected mice had less definition between T cell zones and B cell follicles. To further analyze splenic lymphoid structures, we assessed GC formation during infection by capturing images of individual white pulp regions (Figs 3A and 3B). Initially at D4, infection with either strain led to identifiable GCs in B cell follicles. While Gilliam-infected mice continued to show discernable GCs at D8, Karp-infected mice display scattering of GC B cells within B cell follicles and less apparent GC structures. At D12, the largest contrast was seen between the two strains, as Gilliam-infected spleens maintained clear GCs in B cell follicles whereas Karp-infected spleens presented completely dispersed GC B cells with no identifiable GCs. Overall, our findings establish *Ot*-associated and strain-associated splenic architectural trends. Regardless of the strain or severity of infection, *Ot* infection causes typical splenic architecture to change in lymphoid regions, as we found shrunken white pulp regions in spleens of both infection groups. However, only Karp infection induced GC disorganization, demonstrating GC collapse and GC B cell scattering as unique characteristics.

**Fig 3.**
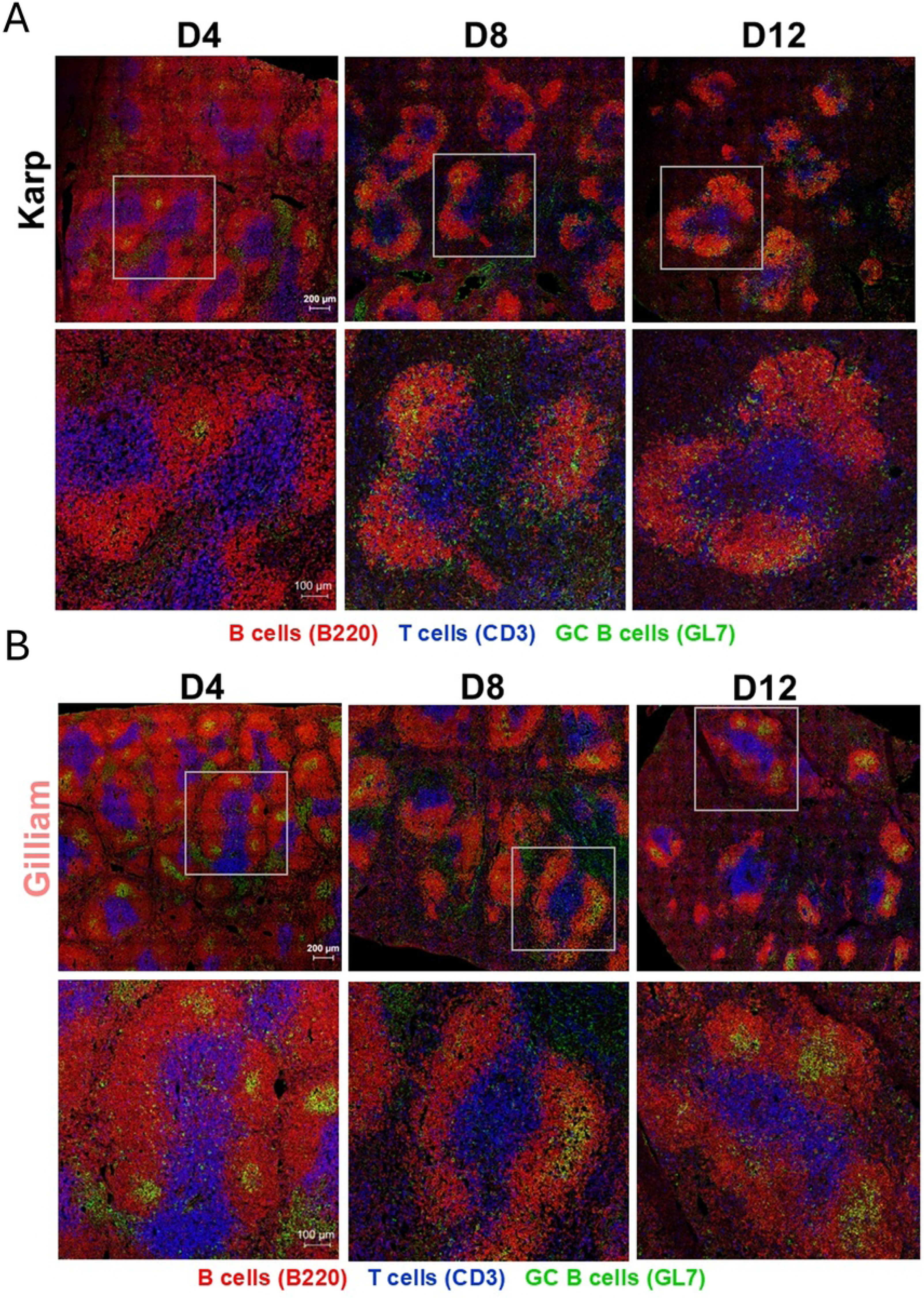
GC disorganization occurs in *Ot* Karp, but not Gilliam infection. Mice were infected, as described in Fig. 1. Cryosections of spleen were prepared and co-stained for B cells (B220), T cells (CD3), and GC B cells (GL7). Images were captured using confocal microscopy (Zeiss LSM 880). Shown are splenic white pulp overview (scale bar, 200 μm) and high magnification images (scale bar, 100 μm) from A) Karp and B) Gilliam infection, respectively.

### Selective impairments of T cell and B cell subsets during Karp, but not Gilliam infection

Splenic GC collapse is linked to the loss of various cellular components in other pathogen infections [37, 38]; however, whether such this mechanism contributes to the altered splenic architecture and GC collapse observed during *Ot* infection remains unclear (Fig 3). By using multi-color flow cytometry (Figs 4A and 4B), we found that the total numbers of T cells, CD4^+^ T cells, CD8^+^ T cells, and activated CD8^+^ T cells were equivalent between Karp- and Gilliam-infected mice, although all of these cell subsets in infected mice were significantly increased as compared to the mock group at various timepoints (Fig 4C and S1 Fig). Importantly, Gilliam-infected mice had significantly higher total numbers of regulatory T cells (Treg) at D8 and D12, as well as increased T follicular regulatory cells (Tfr) at D12 compared to Karp-infected mice. At D12, Gilliam-infected mice had approximately 2.4-fold higher Treg cells and 3.2-fold higher Tfr cell numbers, respectively, than Karp-infected mice, implying greater potential for regulated immune responses during Gilliam infection, but impaired immune regulation during Karp infection.

**Fig 4.**
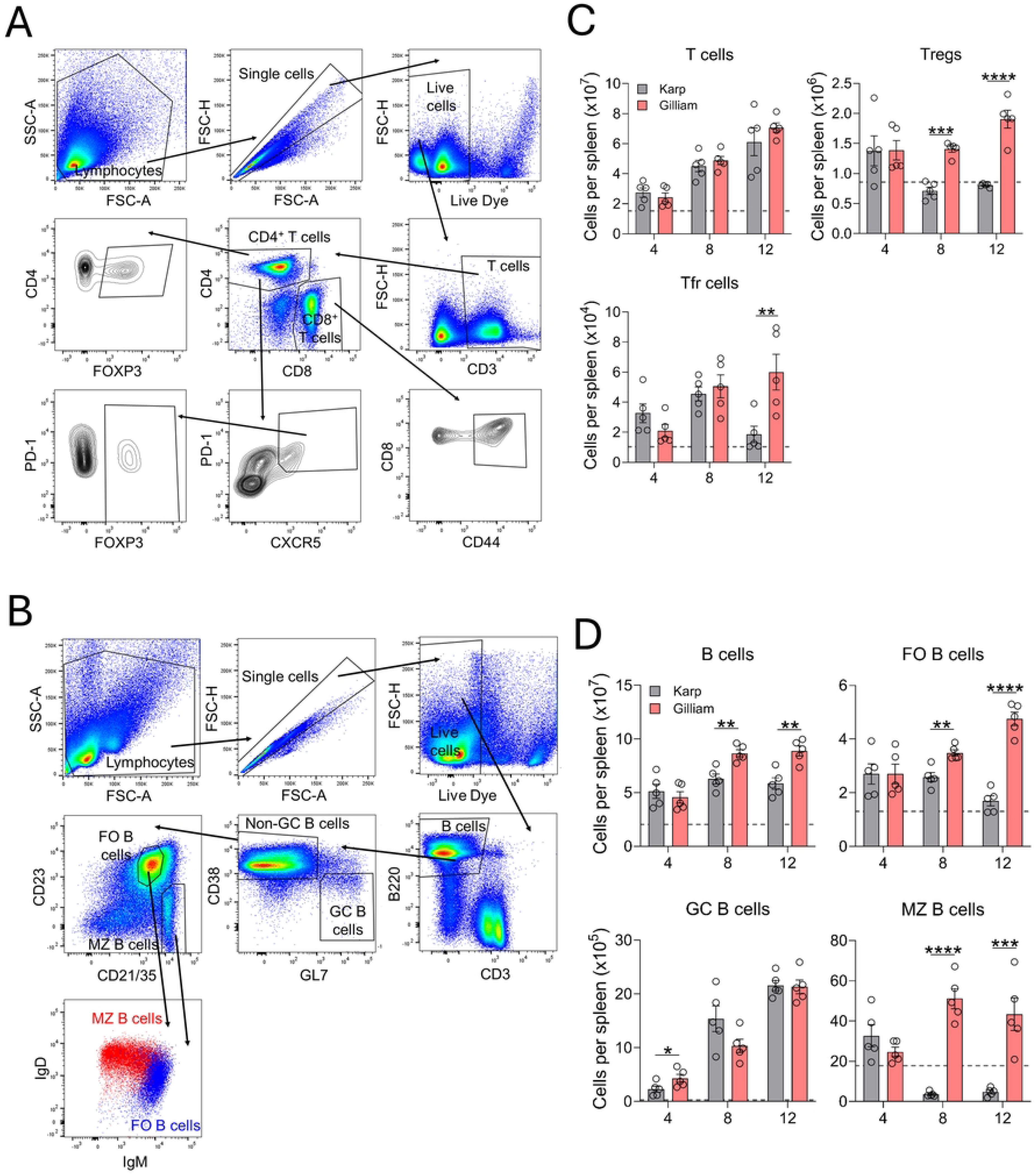
Robust and regulated humoral immune responses elicited by Gilliam, but not Karp infection. Mock and infected mice were euthanized, and their spleens were harvested at given timepoints (5 per group), as described in Fig. 1. Single-cell suspensions were prepared and stained for indicated cell surface markers. A) T cell gating strategy is shown. B) B cell gating strategy is shown. Flow cytometric analysis was performed, and results are presented as C) absolute numbers of various splenic T cell populations; D) absolute numbers of various splenic B cell populations. The absolute cell numbers of the mock group are represented by the dashed grey lines. Data are shown as mean ± SEM from a single experiment and are representative of two independent experiments with similar trends. For statistical analysis, one-way ANOVA was used with a Tukey’s multiple comparisons test for comparison of Karp- and Gilliam-infected samples at each timepoint. Asterisks were representative of comparison between Karp- and Gilliam-infected mice at each timepoint. *, *p* <0.05; **, *p* <0.01; ***, *p* <0.001; ****, *p* <0.0001.

Regarding B cell subsets (Fig 4D), Gilliam infected resulted in a steady increase in absolute numbers of total splenic B, follicular (FO) B, GC B, and marginal zone (MZ) B cell subsets during infection as compared to the mock group (dashed lines). In contrast, Karp-infected mice only showed a significant expansion in GC B cells, notable expansion in total B and FO B cells, but a marked decrease in MZ B cells at D8 and D12 as compared to the mock group. Given these opposite trends, Gilliam-infected mice had nearly 3-fold higher FO B cell numbers at D12, as well as 14-fold and 9-fold MZ B cell numbers at D8 and D12, than Karp-infected counterparts. Collectively, our findings indicate robust and regulated B cell responses during Gilliam infection, but weakened or dysregulated B cell responses at the peak of Karp replication and at severe disease stages.

### Disrupted splenic marginal zones during Karp infection

It is known that systemic *Salmonella* Typhimurium infection can lead to remodeling of splenic microarchitecture, disruption of white pulp topography, and the loss of cell types that compose the MZ, including the reduction in percentage and absolute number of MZ B cells [37]. We and others have reported *Ot* replication in the spleen, implying possible infection of diverse cell types within the MZ (*Ot* is known to infect monocytes, macrophages, dendritic cells, and neutrophils, endothelial cells and fibroblasts) [39–41]. Having demonstrated severe reduction of MZ B cell numbers during Karp infection (Fig 4D), we then used immunohistology to investigate the condition and organization of the splenic MZ during infection. We stained cryosections for B cells (B220), MZ macrophages (SIGNR-1), and metallophilic macrophages (CD169) and generated images via confocal microscopy (Figs 5 A and B). At D4 of both infection groups, the MZ formed a continuous outer layer around B cell follicles with strong, detectable signal for CD169^+^ and SIGNR-1^+^ cells. At D8, alterations in the MZ were noted in Karp-infected spleens, as judged by reduced detection of MZ and metallophilic macrophages, while Gilliam-infected spleens maintained positive signal for CD169^+^ metallophilic macrophages, which formed a dense layer around the white pulp. By D12, positive signal was nearly absent for MZ and metallophilic macrophages, or completely lacked, in some MZ areas of Karp-infected spleens, which was in sharp contrast to Gilliam-infected spleens. Together, these results highlight significant differences in the splenic microarchitecture between Karp-vs. Gilliam-infected mice. Given the role of the splenic MZ in aiding GC responses and orchestrating cytokine and cellular immune responses, the depletion of this lymphoid microstructure in Karp infection may impair not only humoral immune responses, but also cellular immune responses, as suggested in other infection models [42–45].

**Fig 5.**
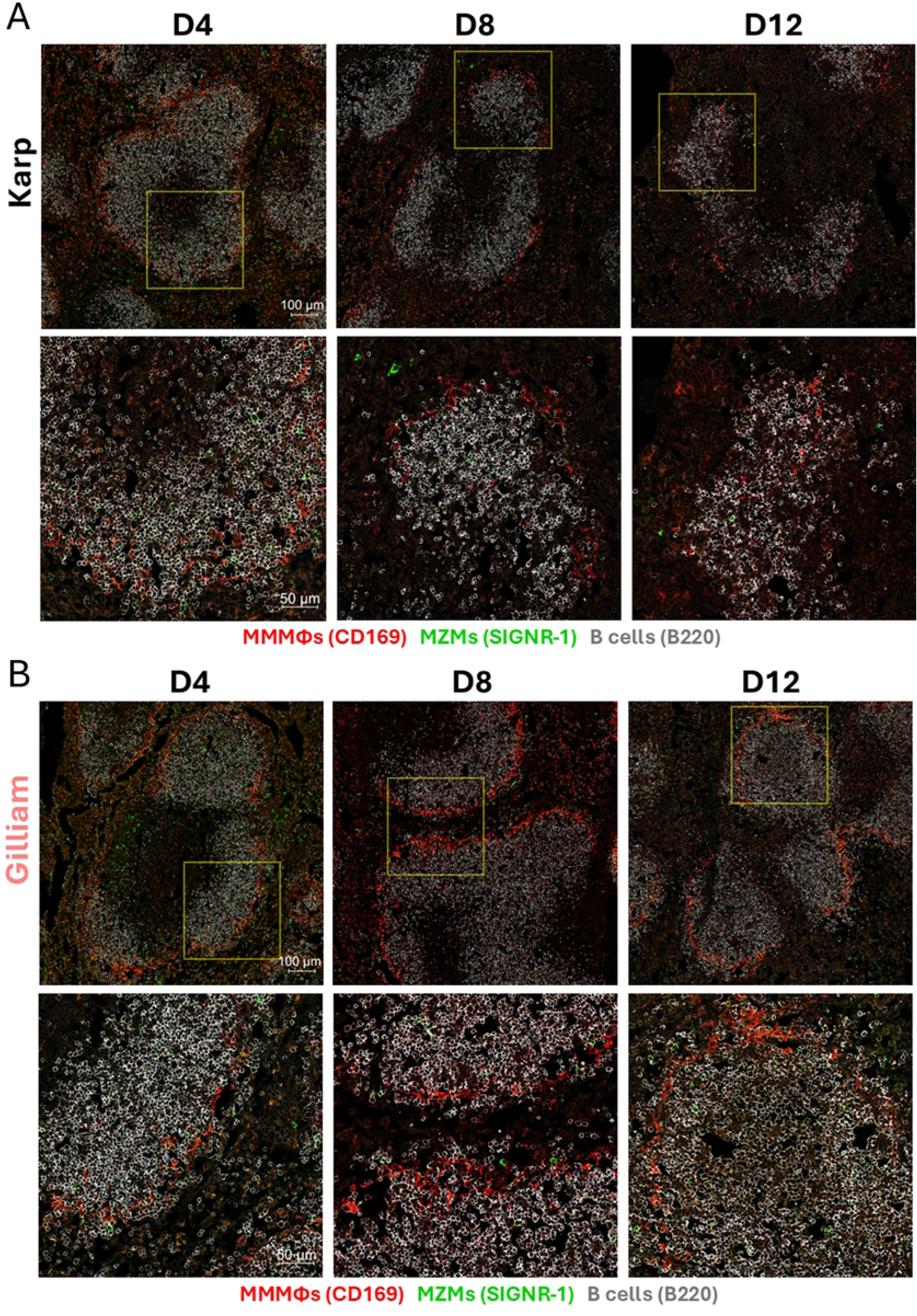
Abrogation of marginal zone occurs exclusively in Karp-infected mice. Mice were infected, as described in Fig. 1. Frozen cryosections of spleen were co-stained for B cells (B220), MZ macrophages (SIGNR-1), and metallophilic macrophages (CD169). Images were captured using confocal microscopy (Zeiss LSM 880) and are shown in overview (scale bar, 100 μm) and high magnification images (scale bar, 50 μm) from A) Karp and B) Gilliam infection are shown.

### Excessive inflammation and activation of leukocyte cell recruitment pathways during Karp infection

GC collapse is related to high inflammatory responses elicited by bacterial infection, implicating regulatory roles for type 1 cytokines (TNFα, IFNγ, IL-12, etc.) [46–48]. Moreover, innate immune cells such as inflammatory monocytes contribute to the loss of GCs [48–50]. To evaluate the status of inflammation and innate immune cells in the splenic microenvironment of Karp- and Gilliam-infected mice, we analyzed spleen samples by using RNAseq. Principal component analysis confirmed that Karp and Gilliam samples clustered distinctively at given timepoints (S2A Fig). Comparison of Karp- vs. Gilliam-infected spleens at D4, D8, and D12 revealed 454, 843, and 1,121 differentially expressed genes, respectively (S2B Fig). To evaluate potential genes encoding biomarkers associated with scrub typhus and inflammation, we plotted log2 fold change of various differentially expressed genes within each group relative to the mock group (Fig 6A). Consistent with our previous reports in murine models [24, 51], Karp infection induced significantly higher expression levels of *Ccl2* (1.8-fold) and *Il33* (1.6-fold) at D4, as well as *Ifng* (2.4-fold) and *Il10* (1.4-fold) at D8. Of note, both IFNγ and IL-10 have been linked to severe scrub typhus in patient-based studies [26]. Additionally, we found that neutrophil degranulation-related genes *Mpo* and *Elane* reached peak expression levels at D12 in both infection groups; however, their expression was higher in Karp infection as compared to Gilliam infection (1.6- and 1.7-fold higher, respectively). We then performed gene set enrichment analysis for a comparison Karp- or Gilliam-infected samples and searched for genes involved with defense and inflammatory responses (Fig 6B). Normalized enrichment scores revealed significant upregulation of defense response pathways involved in “Response to bacterium” in Karp infection at D4. We also identified the upregulation of several inflammatory response pathways during Karp infection, including signaling pathways related to proinflammatory cytokines such as *Il1*, *Il6*, *Il18*, *Ifng*, and *Tnf*, as well as the anti-inflammatory, severe scrub typhus-related cytokine *Il10*. Our analysis also uncovered increased expression of leukocyte recruitment pathways, including granulocytes and monocytes. Together, our results demonstrate a distinctive inflammatory environment in Karp-infected spleens, evidenced by significant upregulation of proinflammatory cytokines characteristic of severe scrub typhus and recruitment of myeloid leukocytes.

**Fig 6.**
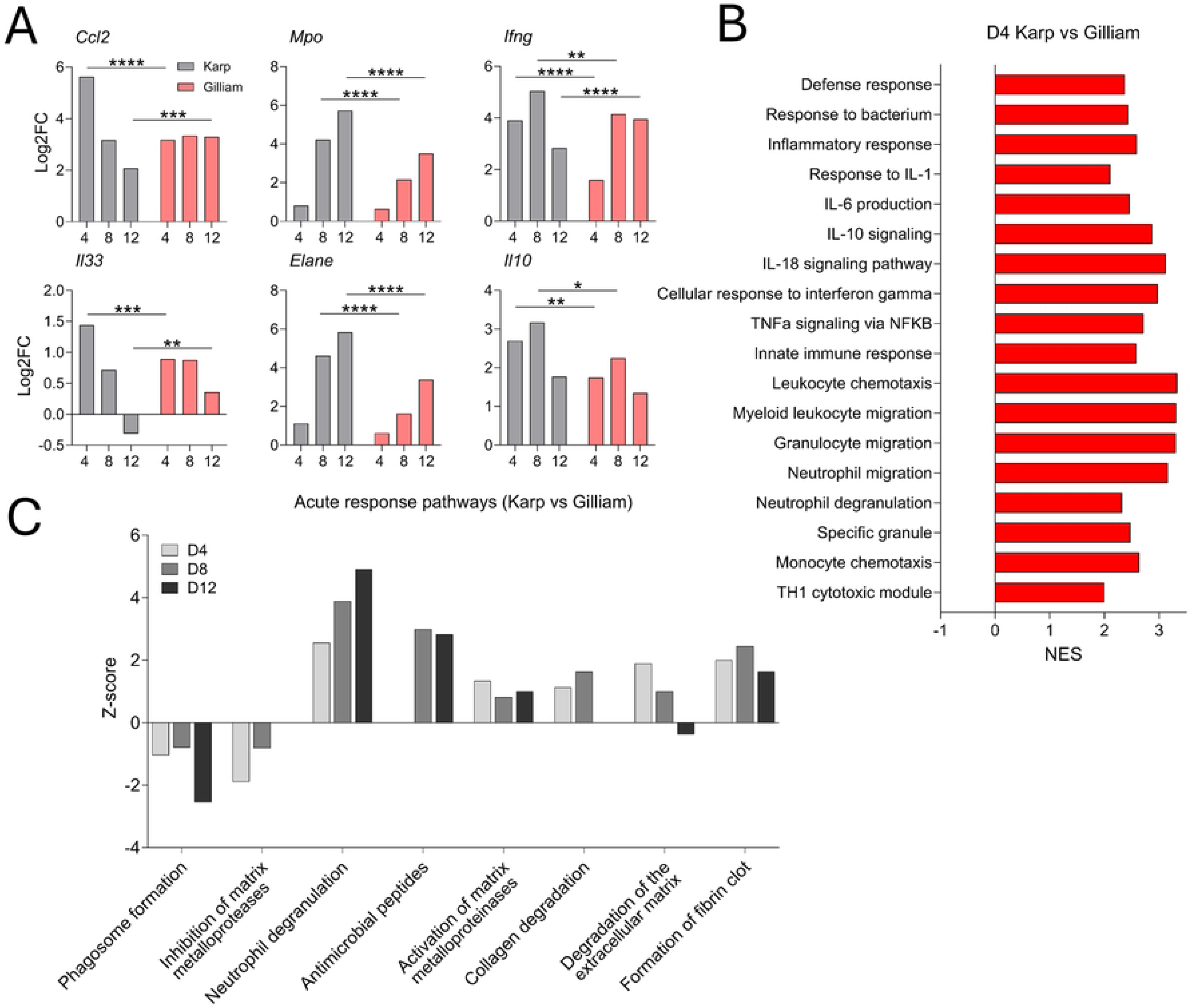
Transcriptomic analysis of splenic environment in Karp- and Gilliam-infected mice. Mice (4 per group) were infected, as described in Fig 1. Spleen tissues were harvested at indicated time-points. RNAseq was performed and analyzed by Novogene, differentially expressed genes from D4 Karp vs Mock, D4 Gilliam vs Mock, D8 Karp vs Mock, D8 Gilliam vs Mock, D12 Karp vs Mock, and D12 Gilliam vs Mock comparisons were identified. A) Select genes are plotted using log2 fold change and displayed. B) Differentially expressed genes relating to inflammatory and innate immune cell pathways were identified by gene set enrichment analysis from D4 Karp vs Gilliam comparison. Pathways upregulated in Karp infection appear as positive NES values. C) IPA analysis of pathways involved in acute phase response from Karp versus Gilliam comparisons at D4, D8, and D12. Results were filtered by a Benjamini-Hochberg corrected p-value. Pathways upregulated in Karp infection show positive z-scores, while pathways upregulated in Gilliam show negative z-scores.

To further characterize the acute response pathways to infection, we used IPA analysis to visualize the expression of pathways from Karp vs. Gilliam comparisons at all timepoints (Fig 6C). Gilliam infection showed upregulation of phagosome formation pathway compared to Karp counterparts during infection, possibly demonstrating increased immune recognition and subsequent phagocytosis. Notably, we found significant upregulation of neutrophil degranulation pathway at all timepoints in Karp infection, which corresponded with the increased expression of pathways capable of extracellular matrix (ECM) remodeling, including activation of matrix metalloproteinases, collagen degradation, and degradation of the ECM. In contrast, Gilliam-infected spleens upregulated the inhibition of matrix metalloproteases pathway at D4 and D8, which could display resistance of ECM degradation in Gilliam infection. These data corresponded with the expression of *Mmp8* and *Mmp9* genes, which showed significantly higher expression in Karp-infected spleens compared to Gilliam counterparts (S3 Fig). We also found significant upregulation of defense response pathways throughout Karp infection, such as formation of fibrin thrombosis and antimicrobial peptides. Next, we assessed the expression of adaptive immune-related pathways between Karp versus Gilliam samples by utilizing IPA analysis at all time points (S4A Fig). Karp infection showed marked upregulation of CTLA4 signaling in cytotoxic T lymphocytes, particularly at D12, with a z-score of 4.747. Yet, Gilliam-infected spleens had increased expression of immunoregulatory interaction pathways at all time-points (z-scores of −1.342 at D4, −0.707 at D8, and −2.53 at D12. Importantly, Gilliam-infected spleens showed upregulation of the lymphoid and non-lymphoid immunoregulatory interactions pathways at all time points compared to Karp counterparts, emphasizing better control of immune responses and corresponding with our observations of expansion of Treg and Tfr cell compartments. At D12, we found elevated expression of Th1 and Th2 pathways in Gilliam infection. Lastly, the dendritic cell maturation pathway showed a unique expression pattern, where Karp-infected spleens demonstrated upregulation of this pathway at D4, but expression of the dendritic cell maturation pathway flipped, and became significantly upregulated in Gilliam counterparts at D8 and D12. Utilizing B cell activation gene lists derived from GO and Biological Process, we identified 33 differentially expressed genes encoding B cell activation components at D12, including *Pax5*, *Cd19*, *Aicda*, and *Cd40* (S4B Fig). Of these genes, 28/33, or 85% were upregulated in Gilliam infection, as compared to only 5/33 genes, or 15% in Karp infection, demonstrating the attenuation of B cell activation in Karp-infected spleens. Our transcriptional analysis uncovered significant differences in the B and T cell response between infection with Karp and Gilliam strains, highlighting the deficiency in adaptive immune signaling pathways in Karp-infected spleens.

### Significant infiltration of activated neutrophils, inflammatory monocytes, and M1 macrophages in Karp-infected spleens

RNAseq analysis of Karp- and Gilliam-infected spleens revealed significant upregulation of myeloid leukocyte migration, including several cell types such as neutrophils and monocytes. To confirm a greater rate of infiltration of myeloid cell subsets in Karp infection, we utilized multi-color flow cytometry to identify several myeloid cell subtypes (Fig 7B). Overall, we found that Karp infection led to greater recruitment of monocytes and inflammatory monocytes (Fig 7B), macrophages and M1 macrophages (Fig 7C), as well as neutrophils and activated neutrophils (Fig 7D). Monocyte (F4/80^+^ Ly6C^hi^) numbers increased during infection with either strain, as compared to mock-infected mice, but were significantly higher in Karp-infected mice compared to Gilliam counterparts at D12. CCR2^+^ SCA-1^+^ monocytes, which have been linked to disruption of GC responses during *Salmonella* infection, were detectable at D4, and showed greater infiltration specifically in Karp infection. By D12, these cells were comparable in Gilliam- and mock-infected spleens, with a 20-fold difference in inflammatory monocyte numbers between Karp and Gilliam infection. Moreover, Karp infection led to a significant increase of total macrophages and M1 macrophages at D12 as compared to Gilliam infection. Given our finding of several upregulated pathways involved in neutrophil migration and degranulation in Karp infection, we assessed the number of neutrophils and activated neutrophils in mock and infected spleens. Our data demonstrates significantly more neutrophils at D8 and D12, as well as activated neutrophils at all timepoints in Karp-infected spleens. Collectively, our myeloid cell flow cytometry data highlights the correlation between significant inflammation and infiltration of myeloid cell subsets in spleens of Karp-infected mice. Thus, uncontrolled recruitment and activation of these myeloid leukocytes may lead to immunopathology and may be linked to GC collapse during Karp infection.

**Fig 7.**
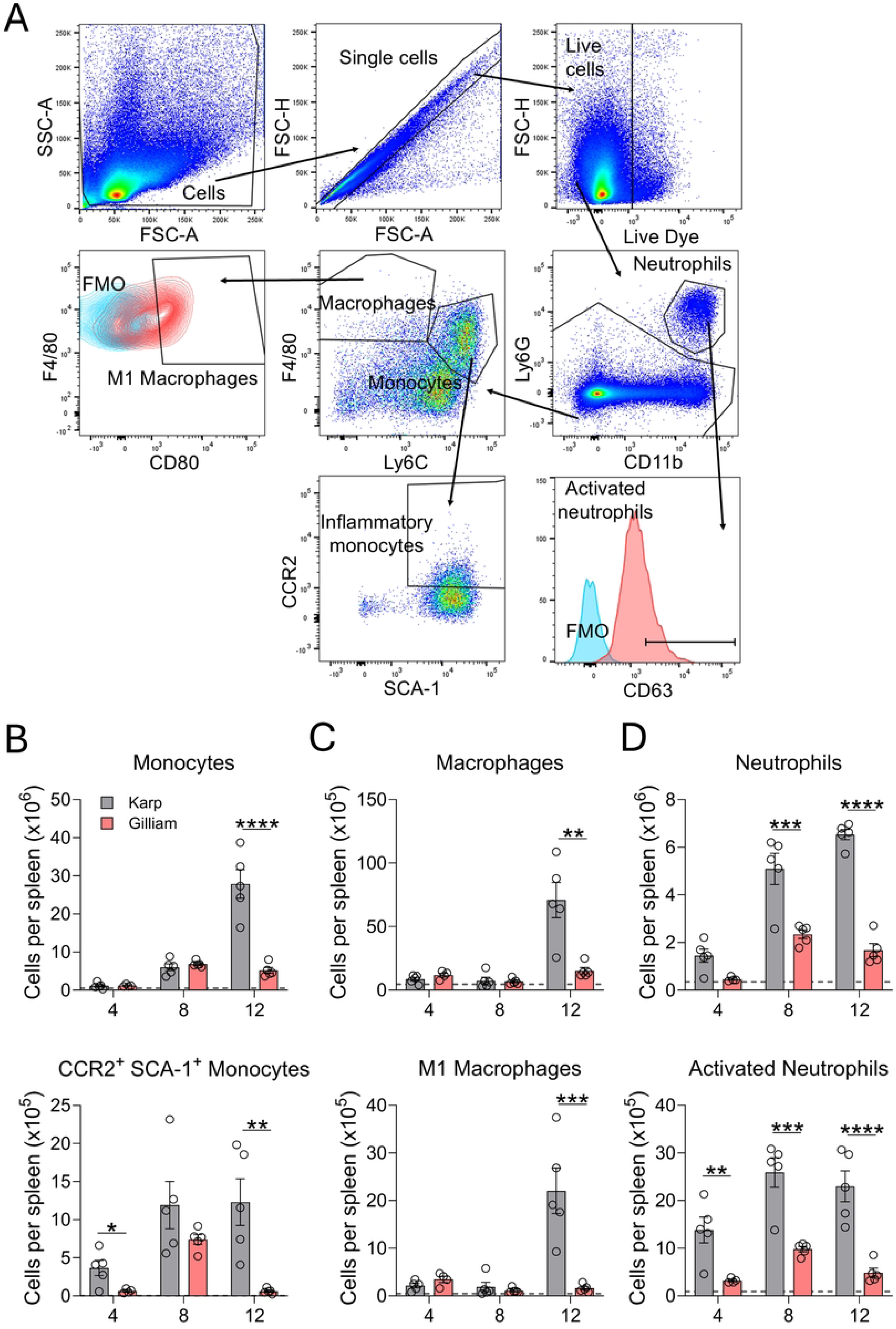
Karp infection induces significant splenic infiltration of myeloid cell subsets. Mock and infected mice were euthanized, and their spleens were harvested at given timepoints (5 per group), as described in Fig. 1. Single-cell suspensions were prepared then stained for indicated cell surface markers. A) Gating strategy for myeloid cell subsets is shown. B) Absolute numbers of monocytes and inflammatory monocytes, C) macrophages and M1 macrophages, and D) neutrophils and activated neutrophils are displayed. The absolute cell numbers of the mock group are represented by the dashed grey line. Data are shown as mean ± SEM from a single experiment and are representative of two independent experiments with similar trends. For statistical analysis, one-way ANOVA was used with a Tukey’s multiple comparisons test for comparison of Karp- and Gilliam-infected samples at each timepoint. Asterisks were representative of comparison between Karp- and Gilliam-infected mice at each timepoint. *, *p* <0.05; **, *p* <0.01; ***, *p* <0.001; ****, *p* <0.0001.

## Discussion

In our previous report, we demonstrated that lethal Karp infection induced disorganization of splenic architecture, impaired humoral immune responses, and loss of GCs [27]. However, it remains unclear as to whether this mechanism is shared across different *Ot* strains that are prevalent in endemic countries. In this study, we provided strong evidence of highly virulent *Ot* strain associated immune patterns such as significant splenic bacteremia and disease scores, GC collapse, excessive inflammation, and increased innate immune cell infiltration in the spleen. Conversely, when mice were infected with a less virulent strain, splenic humoral immune responses were robust, indicated by significantly higher sera IgM and IgG titers during the recovery phase, well-formed GC structures that were maintained throughout infection and greater expression of transcriptomic pathways associated with adaptive immunity. Collectively, these results indicate that severe infection with a highly virulent *Ot* strain results in weakened humoral immunity, particularly GC responses.

Importantly, the collapse of GCs during infection is not unique to *Ot*; in fact, many other pathogens such as *Borrelia burgdorferi*, *Salmonella typhimurium*, *Ehrlichia muris*, SARS-CoV-2, and *Plasmodium* spp have also been proven to induce GC loss [37, 38, 48, 52–54]. Several of these studies have revealed the underlying mechanisms of GC abrogation including recruitment of CCR2-dependent SCA-1^+^ monocytes followed by impaired respiration of GC B cells, loss of follicular dendritic cells, CXCL13 gradient dysregulation induced by excessive TNFα production, and absence of mature Tfh cells [37, 38, 47, 48, 54]. Of note, almost all these mechanisms are associated with uncontrolled inflammatory responses. In the context of *Salmonella* infection, IFNγ induces the differentiation of SCA-1^+^ CCR2^+^ monocytes in the bone marrow. These monocytes migrate to the spleen via CCR2-dependent recruitment, and through an unknown mechanism mediated by TNFα, causing the disruption of GCs [48]. During *Ehrlichia* infection, ablation of TNFα was able to restore organization of splenic white pulp, decrease splenomegaly, and increase splenic GC B cell frequency [47]. Both *Plasmodium* and severe SARS-CoV-2 infections induce diminished GC responses due to the lack of differentiation of mature Tfh cells, a process linked to cytokine storm-like immune responses involving TNFα or a combination of TNFα and IFNγ [38, 54]. Despite the variation in direct mechanisms of GC collapse during infection with a range of pathogens, excessive inflammatory responses are involved in several reports where infection provoked the abrogation of GCs.

Historically, highly virulent *Ot* strains have been associated with more severe disease and excessive inflammation in murine models [24]. Both humans and animal models show unique type 1 inflammatory signatures during severe disease. Recent studies from our lab and others proved that infection with highly virulent strains caused increased gene or protein expression of inflammatory markers in target cells or sera, including the genes *Ccl2*, *Ccl3*, *Ccl4*, and *Tnf* or proteins MCP-1, IL-6, IL-10, and IFNγ [24, 26]. Interestingly, the immune signatures induced by high virulence *Ot* strains contain biomarkers that play a role in infection-induced GC collapse: TNFα, IFNγ, and the major ligand for CCR2, MCP-1. Thus, it is plausible that severe *Ot* infection caused by a high virulence strain initiates similar mechanisms of GC collapse as has been demonstrated with many other pathogens. Indeed, in the splenic transcriptomic environment, we noted significantly elevated expression of *Ifng* and *Ccl2,* as well as the upregulation of pathways involved with *Il1*, *Ill6*, *Tnf*, *Ifng*, and acute inflammatory response. Flow cytometry of infected spleens revealed striking contrast in the infiltration of SCA-1^+^CCR2^+^ monocytes in Karp- and Gilliam-infected spleens. In Karp infection, we identified a significant increase in SCA-1^+^CCR2^+^ monocytes throughout infection, which may disrupt ongoing GC structures that initially formed in the Karp-infected spleens at D4. Interestingly, these inflammatory monocytes were detected and peaked in Gilliam-infected spleens at D8 but were dramatically reduced by 12-fold at D12. Given the GC disrupting role of SCA-1^+^CCR2^+^ monocytes, the persistence of SCA-1^+^CCR2^+^ monocytes in Karp infection but loss of these cells in Gilliam infection at D12, may implicate these inflammatory monocytes in Karp infection-induced GC collapse [48]. Therefore, our data suggest that the type 1 inflammatory immune signatures coupled with infiltrating inflammatory monocytes associated with highly virulent *Ot* strain infection may cause splenic GC collapse.

Beyond inflammatory responses, Karp infection elicited a unique transcriptomic immune signature related to neutrophil responses. Gene set enrichment analysis demonstrated that at D4, Karp infection upregulated pathways including neutrophil migration, neutrophil degranulation, and specific granule compared to Gilliam counterparts. IPA analysis further confirmed the involvement of neutrophils, as pathways encoding neutrophil degranulation and activation of matrix metalloproteinases were significantly upregulated in Karp infection at all time points. In addition, we found that Karp-infected spleens showed significant upregulation of pathways involved in collagen degradation and degradation of the ECM, while Gilliam-infected spleens presented increased expression of genes encoding inhibition of matrix metalloproteases. Flow cytometric analysis of splenic myeloid cells revealed significantly more neutrophil and activated neutrophil absolute cell numbers in the spleens of Karp-infected mice. Of note, neutrophil degranulation, specifically tertiary granules, plays a major role in the remodeling of splenic microarchitecture during *Trypanosoma brucei* infection [55]. The ECM serves as scaffold for cells within the spleen to allow for segregated compartments within the splenic white pulp and the MZ, allowing for organized adaptive immune responses through facilitate cell-cell interactions and cell migration [56]. Upon neutrophil depletion, *T. brucei*-infected mouse survival improved, splenic plasma cell numbers increased, and ECM structure was preserved [55]. Since we uncovered evidence of neutrophil degranulation and degradation of the ECM, it is possible that degranulating neutrophils play a role not only in the collapse of GCs, but also the abrogation of splenic MZs in Karp infection via destruction of ECM noncellular scaffolding.

The structure of the spleen is related to function, where the MZ, composed of MZ B cells, MZ macrophages, metallophilic macrophages, and dendritic cells, establishes the first line of defense against blood-borne pathogens [30]. The splenic MZ is next to the white and red pulp in the spleen, allowing MZ cells to capture blood-borne pathogens and initiate adaptive immune responses in the white pulp, including GC responses [30, 44, 45]. We found a distinctive feature of severe infection caused by the highly virulent Karp strain, the abrogation of splenic MZs. Confocal imaging revealed that macrophage populations of the MZ disappeared, leaving the MZ border nearly absent at D12. Flow cytometry further confirmed that the MZ B cell population was reduced during mid- and late-stage Karp infection. Given the importance of the MZ in GC responses, the abrogation of this region may be a contributing factor in the collapse of GCs during Karp infection. While we have not yet identified the duration of GC disorganization or MZ disappearance, the loss of these structures may impair the development of adaptive immune responses to other infections or even vaccinations [48].

The spleen has a high bacterial burden during *Ot* infection, but how *Ot* infection is established in this organ is unclear [26], and what cell types are its initial targets is not also known [39]. Since the MZ is designed for phagocytic cells to encounter blood-borne pathogens, it is possible that *Ot* may enter the spleen via the blood and establish infection in the phagocytic cells of the MZ. This has been shown for other intracellular pathogens such as *Listeria monocytogenes*, which metallophilic macrophages upon entry into the spleen [57–59]. Given the phagocytic cell tropism of *Ot*, our findings of abrogated splenic MZ warrant further investigation of the MZ as an early target of *Ot* infection.

Changes in splenic white pulp architecture were observed during infection with the Karp or Gilliam strain. We found that white pulp regions were shrunken, and T cells localized outside of T cell zones [37]. This phenomenon has been observed during *Salmonella* infection, where white pulp regions were perturbed even four weeks post-infection [37]. Remarkably, while typical white pulp structure was restored by seven weeks post-infection, MZ regions were still disturbed, with MZ and metallophilic macrophage populations not recovered by this time [37]. Since we observed perturbations in the splenic white pulp regardless of strain or disease severity, it is likely that *Ot*, irrespective of virulence, can alter the microarchitecture of the spleen. While this reorganization diminishes humoral immune responses, as development of B cell responses strongly relies on the structure of the spleen [30]. While a previous study deemed humoral immunity noncritical for *Ot* clearance, it remains unclear whether this abrogation in humoral immunity relates to immune evasion of *Ot* [60]. In fact, reorganized splenic microarchitecture and increased infiltration of innate immune cells may increase encounters of T cells and innate immune cells with the offending pathogen. It is yet to be determined if the disorganization of the splenic microarchitecture is favorable for host by increasing immune cell-*Ot* interactions, or for *Ot* to divert the development of humoral immune responses to persist in the spleen. Similarly, while inflammation can deter the formation of GCs, it is critical for innate and even adaptive immune responses [61]. Overall, our findings suggest that splenic reorganization and inflammation during *Ot* infection may be necessary to clear infection, but it may come at the expense of humoral immune responses and increased immunopathogenesis.

This study has several limitations, some of which warrant further investigation. Firstly, we observed that GC collapse and MZ abrogation occurs in Karp-infected mice, but our experimental timepoints did not allow us to see if these lymphoid structures recover in convalescence or upon resolution of inflammatory stimuli. Future follow-up research with study timepoints weeks or even months post-infection will assess the restoration of these structures, as they play a major role in the development of adaptive immunity. Additionally, the potential use of an attenuated mutant or non-pathogenic *Ot* strain would be helpful to validate some our finding of splenic white pulp disorganization. Since *Ot* remains genetically intractable, our investigation into the influence of *Ot*-carrying phagocytes on lymphoid microarchitecture and GC alterations is limited to use of prototype strains. Future studies with skin inoculation route would help explore B-T cell activation in draining lymph nodes and other lymphoid organs. Of particular interest, we identified the presence of early serum IgG responses in Karp infection, which may contribute to the proinflammatory environment in Karp-infected spleens through FCγR functions on leukocytes [62]. Beyond driving inflammatory responses, early IgG responses in Karp infection may have the capacity to induce antibody dependent *Ot* growth enhancement, as prior data have supported this notion [63, 64]. Future experiments to address the role of antibodies during infection, whether immunopathogenic, autoreactive, or protective, in the context of *Ot* infection with strains of differential virulence will shed light on the function of humoral immune responses in diverse scrub typhus disease models. Lastly, a breakthrough future study would address how inflammation (or given molecules) contributes to GC collapse and lymphoid architecture disruption during severe *Ot* infection using genetic knockout or knock-in mouse models.

In summary, this study uncovered an *Ot* virulence-associated trend of impaired GC and humoral immune responses during severe infection, which may be linked to excessive inflammation and innate immune cell infiltration. Karp infection induced higher splenic bacterial burden, neutrophil infiltration, MZ perturbation, and increased expression of inflammatory genes and related pathways. On the other hand, Gilliam infection caused significantly lower bacterial burden, higher sera antibody responses at late-stage infection, and elevated expression of adaptive immune response pathways compared to Karp counterparts. Yet, regardless of the severity of infection or *Ot* strain virulence, we noted altered splenic white pulp architecture and T cells scattered outside of T cell zones. Our study has provided unique insights into the abrogation of humoral immune responses during severe *Ot* infection and showed evidence of an inflammation related mechanism that has been demonstrated to provoke GC collapse and disruption of splenic microarchitecture in other infections. This study helps fill the knowledge gap as to how *Ot* strain virulence influences the acute adaptive immune response to infection.

## Acknowledgements

We would like to thank the UTMB Flow Cytometry & Cell Sorting Core Lab (Meredith Weglarz), the Optical Microscopy Core for our data acquisition and analyses, and Biostatistics Core (Drs. Xiaoying Yu and Yuanyi Zhang) for RNAseq data analysis. We also thank Dr. David Walker, Dr. Robert Abbott, Nicole Mendell, Layne Pruitt, and Nicole Weidner for their contributions to this project.

## Supporting Information

**S1 Table. Primer sequences for RT-PCR analysis used within the study.**

**S1 Fig. Equivalent absolute cell numbers of T cell subsets in infected spleens.** Mock and infected mice were euthanized, and their spleens were harvested at indicated timepoints (n = 5 per group), as described in Fig. 1. Single-cell suspensions were prepared and stained for indicated cell surface markers and gated, as described in Fig. 4. Flow cytometric analysis was performed, and results were presented as absolute numbers of various splenic T cell populations. Data are shown as mean ± SEM from a single experiment and are representative of two independent experiments with similar trends. For statistical analysis, one-way ANOVA was used with a Tukey’s multiple comparisons test for comparison of Karp- and Gilliam-infected samples at each timepoint. Asterisks were representative of comparison between Karp- and Gilliam-infected mice at each timepoint. *, *p* <0.05; **, *p* <0.01; ***, *p* <0.001; ****, *p* <0.0001.

**S2 Fig. Overview of differentially expressed genes identified by RNAseq Karp vs. Gilliam comparisons.** Mice were infected, as described in Fig. 1; RNA was isolated from spleen tissues of infected mice and sent for RNAseq analysis by Novogene (n = 4 per group). Data were normalized and differentially expressed genes were identified by using the DESeq2 R library. Principal Component Analysis was performed on the FPKM expression values (A). Differentially expressed genes detected by Karp versus Gilliam comparisons were done at D4, D8, and D12 on all samples and plotted (B).

**S3 Fig. B cell activation differentially expressed genes from RNAseq Karp vs. Gilliam comparison at D12.** Mice were infected, as described in Figure 1; RNA was isolated from spleen tissues of infected mice and sent for RNAseq analysis by Novogene (n = 4 per group). Data were normalized and differentially expressed genes were identified using DESeq2 R library. B cell activation gene list was downloaded from Gene Ontology and Biological Process. Differentially expressed genes from B cell activation gene list were plotted and shown in heatmap from Karp vs. Gilliam comparison at D12 (A).

**S4 Figure-*Mmp8* and *Mmp9* differential expression from RNAseq.** Mice were infected, as described in Figure 1; RNA was isolated from spleen tissues of infected mice and sent for RNAseq analysis by Novogene (n = 4 per group). Differentially expressed genes from D4 Karp vs. Mock, D4 Gilliam vs. Mock, D8 Karp vs. Mock, D8 Gilliam vs. Mock, D12 Karp vs. Mock, and D12 Gilliam vs. Mock comparisons were identified. Select matrix metalloprotease-encoding genes were plotted using log2-fold change and displayed.

